# Estimating the rate of quantitative trait evolution in the presence of gene tree discordance by calculating likelihoods across trees

**DOI:** 10.1101/2025.10.06.680724

**Authors:** Yu K. Mo, Matthew W. Hahn

## Abstract

Quantitative traits provide insights into how phenotypes evolve across species. However, standard comparative methods often assume a single species tree and overlook the discordant gene tree histories that may underlie complex traits. Here, we develop a model and software (Spinney) that explicitly incorporate gene tree heterogeneity into rate estimation. Spinney finds the optimal rate of evolution and ancestral states by jointly maximizing the likelihoods across a set of gene trees. Using simulated data, we compare rate estimates from Spinney with those using the species-tree alone, test Spinney’s ability to distinguish true rate variation from spurious signals caused by gene tree discordance, and evaluate ancestral state reconstruction. Spinney consistently produced more accurate rate estimates and reduced incorrect inferences of rate variation. This method therefore provides a flexible framework to integrate gene tree heterogeneity into comparative methods and to produce reliable inferences of quantitative trait evolution, regardless of the source of discordance.

## Introduction

The rate at which quantitative traits evolve among species can tell us much about the evolutionary forces affecting phenotypes. Previous studies have used quantitative traits to investigate the strength and frequency of adaptive evolution (e.g. Hansen 1997; Khabbazian et al. 2016), the effect of changes in selective regimes on trait evolution (e.g. Rafferty and Nabity 2017; Bertram et al. 2023), and to understand the traits underlying species diversification (e.g. Uyeda et al. 2011; Helmstetter et al. 2023). Phylogenetic comparative methods (PCMs) are widely applied to make such inferences on trees (e.g. Felsenstein 1985; Pagel 1999; Blomberg et al. 2003; O’Meara et al. 2006; Adams 2014).

Most PCMs make the fundamental assumption that the species tree represents the evolutionary history of traits. While there may be issues if the species tree is inferred in error (Martins 1996), a larger problem can occur when a trait’s actual history does not match the species history. Gene tree discordance describes topological differences both among individual gene trees and with the species tree. Discordance is ubiquitous among eukaryotes, largely due to incomplete lineage sorting (ILS) and introgression (Degnan and Rosenberg 2009). Because quantitative traits are determined by genes that may be discordant, such traits do not have to have the same history as the species tree, leading to inaccurate inferences. For example, evolutionary rates are consistently overestimated in the presence of even low levels of discordance due to ILS (Mendes et al. 2018). Similarly, Adams et al. (2025) demonstrated that phylogenetic regression models inaccurately estimate trait-environment relationships when there is gene tree discordance due to ILS. Discordance due to introgression can also lead to inaccurate inferences when one assumes that traits follow a bifurcating species tree (e.g. Bastide et al. 2018; Hibbins and Hahn 2021).

Multiple studies have attempted to model traits that may have discordant histories. Bastide et al. (2018) allowed quantitative traits to evolve along both the species tree and an introgression history. While this does not model gene tree discordance directly, it does allow traits to have multiple histories. More recently, Hibbins et al. (2023) developed two methods that allowed for discordance due to ILS or introgression, showing that both led to more accurate inferences. The first method incorporated gene tree discordance into the construction of the covariance matrix, *C*, used by many PCMs. The second method used the pruning algorithm (Felsenstein 1973) to calculate the weighted likelihood of quantitative trait evolution over the tree topologies possible in a species tree. Although these latter results were limited to three species, the pruning algorithm offers many advantages because of its ability to specify diverse transition matrices, to efficiently calculate likelihoods, and to reconstruct ancestral states (Freckleton 2012).

Here, we introduce a new method for inferring evolutionary rates and ancestral states of quantitative traits. This new method—implemented in the program, Spinney—is built on the pruning algorithm framework of Hibbins et al. (2023), but accommodates any number of species. In addition, Spinney introduces new methods both for accurately estimating ancestral states and for accommodating multiple evolutionary rate regimes across a species tree in the presence of discordance. We demonstrate the utility and accuracy of Spinney using simulated data, finding the areas of tree space where it is most useful.

### New Approaches

#### Rate estimates inferred from a set of gene trees

Our method models a continuous trait evolving via Brownian motion. The transition matrix of this model is determined by the probability density function of Brownian motion, where *x*_*i*_ and *x*_*f*_ represent the initial and final trait values over time interval *t*, and *σ*^2^ is the evolutionary rate parameter:

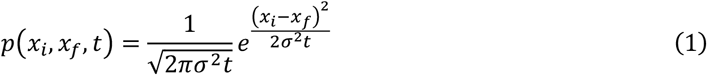

To infer the rate parameter, *σ*^2^, while accounting for discordance, our method uses a group of gene trees (a “spinney”) with branch lengths in coalescent units, along with weights for each gene tree (*w*_*i*_; Figure 1). If each of *n* trees is unique, the weight of each is simply 1/*n*. The likelihood of each gene tree given data at the tips is computed individually using Felsenstein’s pruning algorithm (the same data is used for each tree). Spinney calculates the likelihood for each tree in parallel; this reduces runtime, especially when *n* is large. The total likelihood is obtained by summing the weighted likelihoods of individual gene trees, which is then used for numerical optimization to infer *σ* ^2^ and ancestral states:

**Figure 1.**
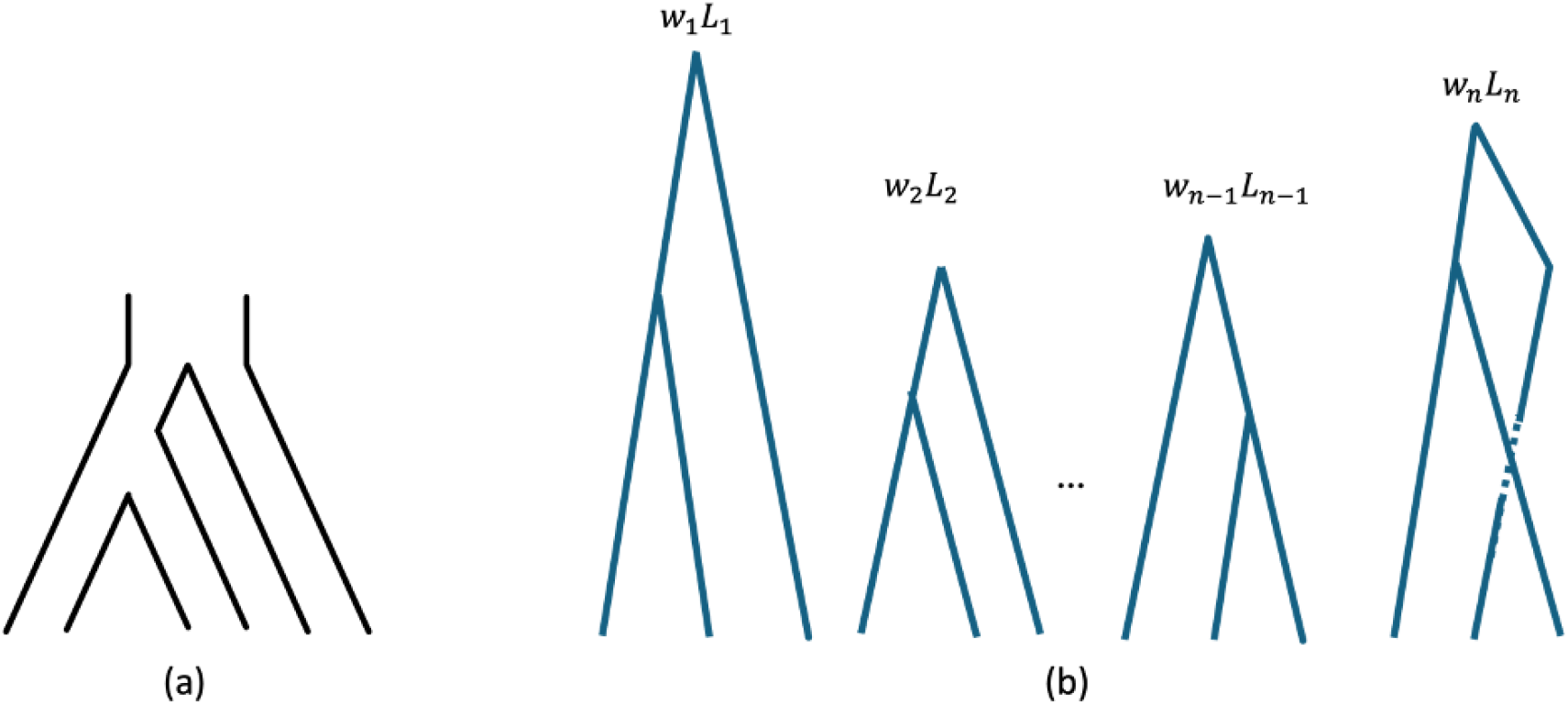
Spinney uses a set of trees for inference. Given (a) a species tree, (b) a set of gene trees is sampled. For *i*-th gene tree, the weight is denoted as *w*_*i*_ and the likelihood is *L*_*i*_. Inferences are carried out based on the summation of weighted likelihood across the gene trees.

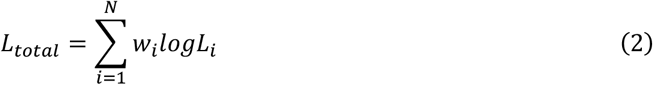

As in Hibbins et al. (2023), our model can take as input ultrametric gene trees either estimated from data or sampled from the species tree under the multispecies coalescent (MSC) model (or any other arbitrary model). We do not assume that we have sampled gene trees at the exact loci underlying our quantitative trait, only that our sample is representative of such trees. Spinney is implemented in Python using several standard libraries. NumPy (Harris et al. 2020) provides basic numerical calculations and DendroPy (Moreno et al. 2024) supports tree processing. Rate optimization is carried out with the Nelder–Mead algorithm from SciPy (Vertanen et al. 2020), and parallelization is implemented with multiprocessing. The source code is freely available on GitHub (https://github.com/speechlesso/spinney).

#### Incorporating rate variation using branch segments

The rate of evolution of quantitative traits can vary across a phylogeny, with the *σ* ^2^ parameter being higher or lower in different clades. One important use of PCMs is in identifying this rate variation (O’Meara et al. 2006). However, because more discordance can result in higher rate estimates, some parts of the species tree may give false signals of rate acceleration simply because of different levels of gene tree heterogeneity (Hibbins et al. 2023). Here, our aim is to accommodate different levels of discordance when testing for the presence of multiple rate-parameters across a species tree.

When calculating shifts in rates directly on a species tree, rates are naturally set to change at speciation nodes. However, when modeling traits on gene trees, internal nodes of such trees do not correspond to speciation events. Furthermore, these internal nodes do not have to be the same age across gene trees. To deal with this heterogeneity, we introduce the idea of unary “time-slicing” nodes that correspond to the age of speciation events (Figure 2a). In this approach, evolutionary rates (i.e. *σ*^2^ parameters) are assigned to branches of the species tree; each gene tree contains time-slicing nodes at times corresponding to speciation events, provided that the lineage at that time has the same set of descendants as in the species tree (cf. Ogilvie et al. 2017; note that this does not imply that gene trees must be concordant). These nodes segment each gene tree, ensuring that all segments corresponding to the same branch in the species tree share the same rate (Figure 2b, 2c). In this way, rates occurring on different branches of the species tree can correctly be applied to the appropriate branches of each gene tree.

**Figure 2.**
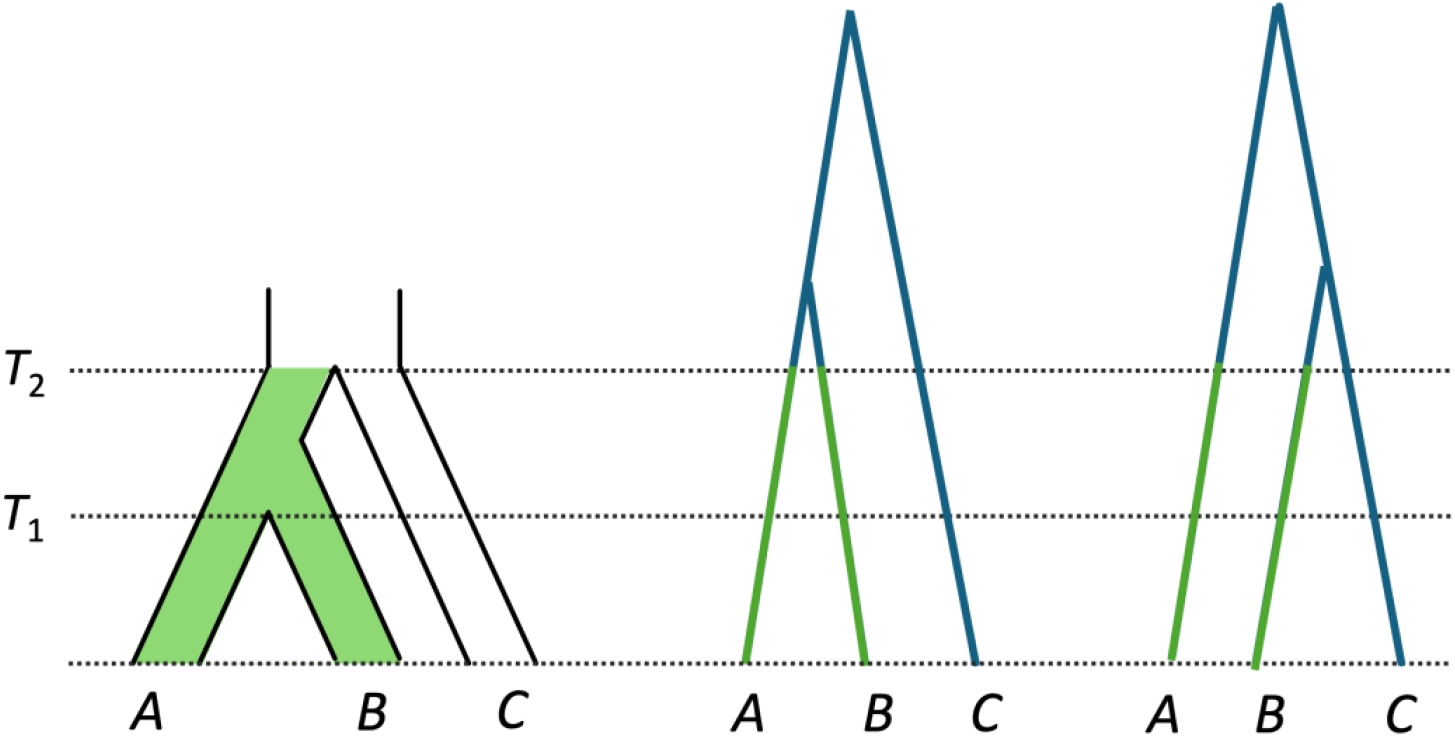
Spinney allows multiple rates within a gene tree. (a) Given a species tree with speciation time *T*_1_ and *T*_2_, evolutionary rates are assigned to branches of this tree. Here we imagine that there are two rates in the species tree (green and white branches). Gene tree segments that exist within the same species tree branch share the same evolutionary rate, whether a gene tree is (b) concordant or (c) discordant.

#### Ancestral state reconstruction

Reconstructing ancestral states traces how traits change through evolutionary history, revealing when shifts occurred and whether similar traits evolved independently in different lineages. However, a challenge again arises because gene trees may differ in both topology and internal-node ages from species trees.

To resolve this problem, we again use the same time-slicing nodes described above. These nodes provide a unified timeframe, allowing discordant gene trees to be combined and ancestral states to be mapped back onto the species tree. Ancestral states are typically estimated at speciation nodes of a phylogeny (e.g. Schluter et al. 1997), even though gene trees do not coalesce until after speciation (looking backward in time; Gillespie and Langley 1979). Time-slicing nodes allow us to obtain maximum likelihood estimates of ancestral states at speciation times averaged across individual gene trees. For a given speciation event, we identify taxa associated with the event, identify corresponding branches from relevant gene trees, and add time-slicing nodes to those branches. The ancestral state is determined by weighting the trait vectors recorded for each gene tree according to tree weights.

## Methods

### Simulating traits and gene trees

We conducted a series of simulations to evaluate the performance of our method. We simulated 1000 gene trees with ms (Hudson 2002) under each species tree described below. We also constructed a covariance matrix, *C*^*^, from the simulated gene trees using the R package seastaR, in order to incorporate discordant histories (Hibbins et al. 2023). For each condition, we then simulated 100 independent traits under a multivariate normal distribution. The trait values across species were drawn as **X** ~𝒩(**0**, Σ⨀*C*^*^), where **0** is a zero vector for the ancestral values, *C*^*^ is the covariance matrix derived from simulated gene trees, and Σ is a scaling matrix of the same dimension as *C*^*^. Σ controls the rate structure of trait evolution. When Σ = *σ*^2^(**11**^*T*^), it corresponds to a scenario with a homogeneous rate *σ*^2^. More generally, it can take different values across entries, allowing simulation with heterogeneous rates across branches of the species tree.

Gene trees used for likelihood calculations were also sampled from ms based on a given species tree; this is equivalent to assuming the MSC model with no introgression. To balance computational efficiency with inference accuracy, we limited the number of sampled gene trees to a sufficiently large yet manageable number (Supplementary Figure 1). We used 20 gene trees as the default for inference, where each tree has the same weight, unless otherwise specified. We use 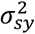 and 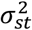 to denote estimated rate parameters inferred by Spinney and the species tree, respectively.

### Single-rate tests

For the single-rate tests, we started with a four-taxon tree (Figure 3a), keeping the total tree height fixed at 10 coalescent units while adjusting internal branch lengths to obtain discordance levels of 1.6%, 25%, and 50%. To test how Spinney generalizes to larger trees, we added cases with 5, 10, 20, and 50 taxa. The species trees for these larger trees were simulated using the R package ape (Paradis and Schliep 2019), with birth and death rates set to 0.1. The discordance levels among these trees are 53%, 66%, 68% and 98%, respectively. The evolutionary rate was kept constant (*σ*^2^ = 1) for all single-rate tests.

**Figure 3.**
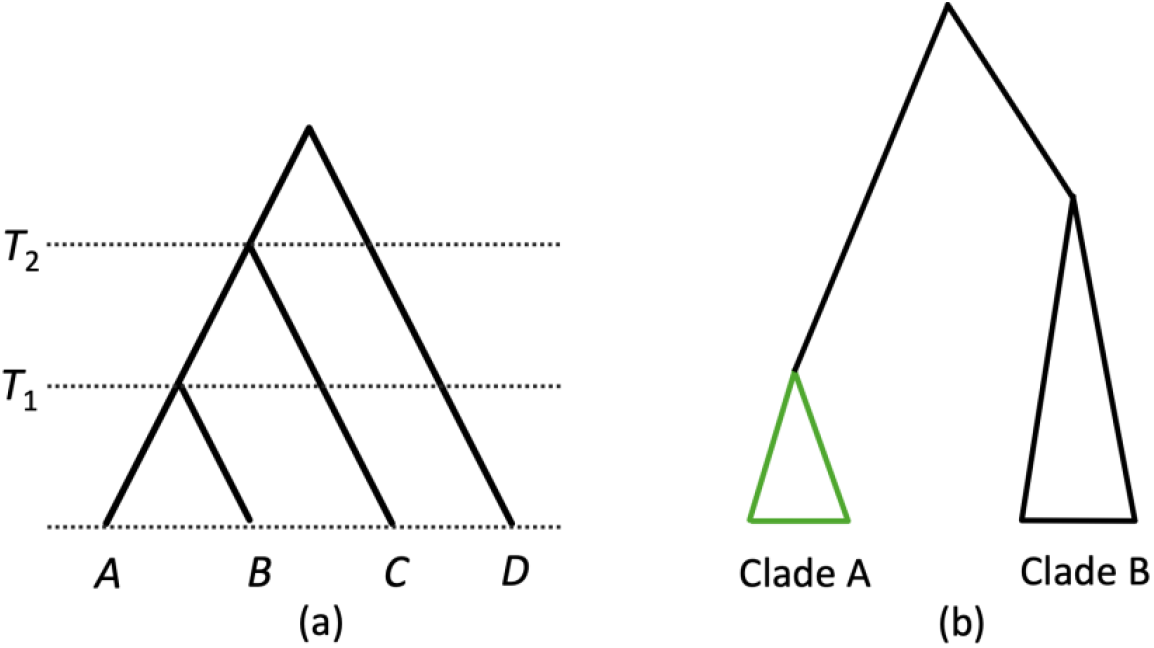
Species trees used for experiments. (a) Four-taxon species tree. Traits are simulated with a single, uniform rate. Speciation times are noted. (b) Two-clade species tree. Traits are simulated either with a single, uniform rate (to test the false positive rate) or with two rates (to test the true positive rate). In all simulations and inferences with two rates, the branches labeled in green have a different rate from the branches labeled in black. Within the species tree, clade A and clade B have different levels of discordance where clade A contributes all discordance and clade B has no discordance. Detailed trees with branch lengths are shown in Supplementary Figure 2.

### Two-rate tests

We used several scenarios to evaluate methods for detecting multiple rate parameters. We started with a species tree with clades A and B (Figure 3b, Supplementary Figure 2a). To ensure no discordance between clades, we extended the internal branches of both clades sufficiently. Consequently, almost all discordance occurs in clade A, leading to 50% discordance. Singe-rate simulations were run with *σ*^2^ = 1, and additional scenarios were tested by modifying *σ*^2^ to 0.1 and 10 with the same species tree. Two-rate simulations were conducted on a different tree (Supplementary Figure 2b) by having 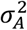 (the evolutionary rate in clade A) take on the values 0.01, 0.1, 1, 10, and 100, while keeping the rest of the tree at *σ*^2^ = 1.

To estimate the false positive rate, we estimated models with both one value and two values of *σ*^2^, using both Spinney and calculations directly on the species tree (all data were produced using simulations with only one rate parameter in this setting). Models with two-rates were fit so that clade A was assigned a distinct rate parameter from the rest of the tree (this is the clade that contains most of the discordance). We used a likelihood ratio test (LRT) to assess the false positive rate of both approaches: any result that favors two rate-parameters is taken to be a false positive. Given a trait and a set of gene trees, we computed the likelihoods under both one-rate (*L*_1_) and two-rate (*L*_2_) models. The test statistic was computed as:

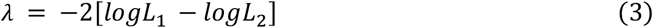

followed by a χ^2^ test with one degree of freedom and a significance threshold of 5%.

### Evaluating ancestral states

We examined how ancestral state reconstruction is affected by using either the species tree alone or Spinney. To evaluate this, we used the same four-taxon simulations as in the single-rate tests. Rates were inferred using a one-rate model under both the species tree and Spinney, and ancestral states were taken as the estimates associated with the MLE of *σ*^2^. Our primary focus was on the speciation event that separates species *A* and *B* from species *C* (at time *T*_2_; Figure 3a).

To visualize ancestral state changes over time, we divided the tree into 100 equally wide bins of time. At each time slice (i.e. bin boundary), we recorded the estimated ancestral state. The ancestral state at the time slice immediately following a speciation event was used as an approximation of the state at that speciation time (because speciation events may not correspond exactly to the boundary of two bins). The ancestral state is represented internally as a vector of the same length as the discretized trait vector; each entry in this vector indicates the probability of observing a specific trait value. Additionally, we computed the 95% confidence interval (CI) for the ancestral state. To determine the CI, we calculated the cumulative distribution function (CDF) from the ancestral state vector. The lower bound for trait *x*_*i*_ was identified as the point where CDF(*x*_*i*_) ≥ 0.025 and the upper bound for trait *x*_*j*_ was where CDF(*x*_*j*_) ≥ 0.975. The width of the confidence interval was then calculated as *x*_*j*_ − *x*_*i*_.

## Results

### Spinney mitigates rate overestimation

For the single-rate case, we compared rate estimates obtained using Spinney with those inferred directly from the species tree. All calculations on the species tree use the standard pruning algorithm to optimize *σ*^2^ on a tree with correctly specified species tree branch lengths. Our primary goal was to assess whether Spinney could accurately estimate the rate of evolution while accounting for gene tree heterogeneity (here caused by ILS), as opposed to using the species tree, which does not incorporate discordance.

For four-taxon trees, inference using the species tree resulted in increasingly severe rate overestimation as discordance increased (Figure 4a). Across 100 replicates, the mean 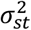 values were 1.44, 3.66, and 12.47 for discordance levels of 1.6%, 25%, and 50%, respectively (the true value of *σ*^2^ is always 1). Notably, even a small level of discordance led to overestimation. In contrast, Spinney produced more accurate rate estimates. The mean 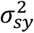 values were 1.03, 1.60, and 3.30 for discordance levels of 1.6%, 25%, and 50%, respectively. While higher levels of discordance still led to overestimation of rates, this bias was mitigated by using Spinney. Additionally, the standard deviation of rate estimates from Spinney was smaller compared to those from the species tree (Supplementary Table 1).

**Figure 4.**
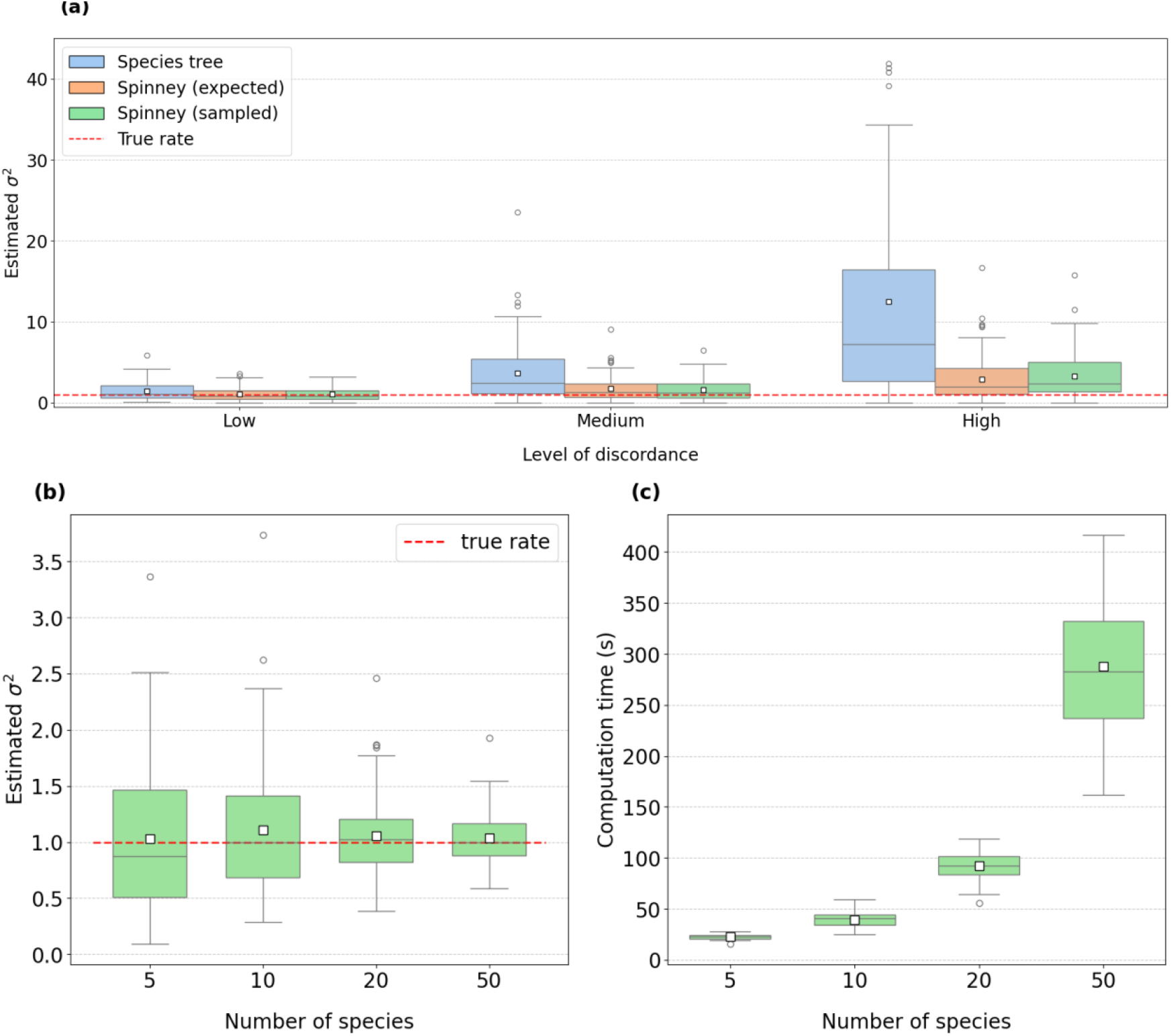
Rate estimates for the single-rate case. Rates were inferred using the species tree and Spinney. (a) Four-taxon settings under discordance levels of 1.6% (Low), 25% (Medium), and 50% (High). Spinney used expected gene trees derived from the MSC (orange boxes) and sampled gene trees (green boxes). (b) Rates were inferred by Spinney for species trees with 5, 10, 20, and 50 taxa, with 53%, 66%, 68% and 98% discordance, respectively. Spinney only used sampled gene trees in these calculations. (c) Computation time for the analyses in panel b. Runtime increases with the number of species.

In the above results, Spinney was run by calculating the partial likelihoods across a set of 20 gene trees. We also ran Spinney with only four gene trees: those expected under the MSC model for each discordance level. In our setting, only the branch subtending species *A* and *B* has discordance, so there are only four gene trees, with weights determined by the MSC model. These expected sets are equivalent to a sufficiently large set of sampled gene trees (Supplementary Figure 1). Rate estimates from expected gene trees closely matched those from sampled gene trees, consistently correcting overestimation. These also showed lower standard deviations and fewer outliers, as with sampled gene trees (Figure 4a).

Generalization to larger species trees (5, 10, 20, and 50 taxa) revealed the same pattern: Spinney corrected overestimation relative to species-tree rates (Figure 4b). For example, for the 10-taxon case, the mean rate estimates by the species tree and Spinney are 2.00 and 1.11 respectively. Although discordance levels frequently exceeded 50% in these larger trees, the degree of overestimation was less severe than in the four-taxon scenario.

Unlike in the four-taxon trees, it is generally infeasible to enumerate all possible topologies under the MSC model for larger trees, as the number of topologies increases double-factorially with the number of taxa (Felsenstein 2004). Spinney must therefore rely on a finite sample of gene trees; but, even with moderate sample sizes (e.g., 20 gene trees), it produced accurate and stable estimates.

We recorded the computer wall-clock time for large trees. As expected, runtimes increased with the number of species. For 5, 10, and 20 taxa, each replicate finished in 28, 98, and 119 seconds respectively (Figure 4c). For 50 taxa, wall-clock time ranged from 162 to 417 seconds. All optimizations ran on Indiana University’s Big Red 200 (HPE Cray EX, SLES 15) CPU nodes (AMD EPYC 7742). Spinney used one node per condition, with 10 threads and 2 GB memory.

### Spinney reduces false inferences of rate variation

We are particularly interested in whether discordance could lead to incorrectly favoring a model with two rate parameters when there is more discordance in one part of a tree. Specifically, we predict that calculations using the species tree would favor the two-rate model because high discordance in (for instance) clade A will falsely leads to a distinct *σ*^2^ compared to the rest of the tree (Figure 3b). We also predict that Spinney will correct this overestimation by integrating discordant histories, thereby reducing the tendency to favor the two-rate model.

As expected, inference by the species tree overestimated rates. Starting with the one-rate model applied to the tree in Figure 3b, the mean 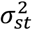 was 8.56, whereas the mean 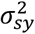 was 1.93. When estimating a two-rate model (only one rate was used to simulate these data), both methods inferred a higher rate for clade A and recovered an accurate rate for clade B. However, the species tree overestimated 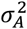 by 18X, while Spinney was only 5X higher (Figure 5; Supplementary Table 3).

**Figure 5.**
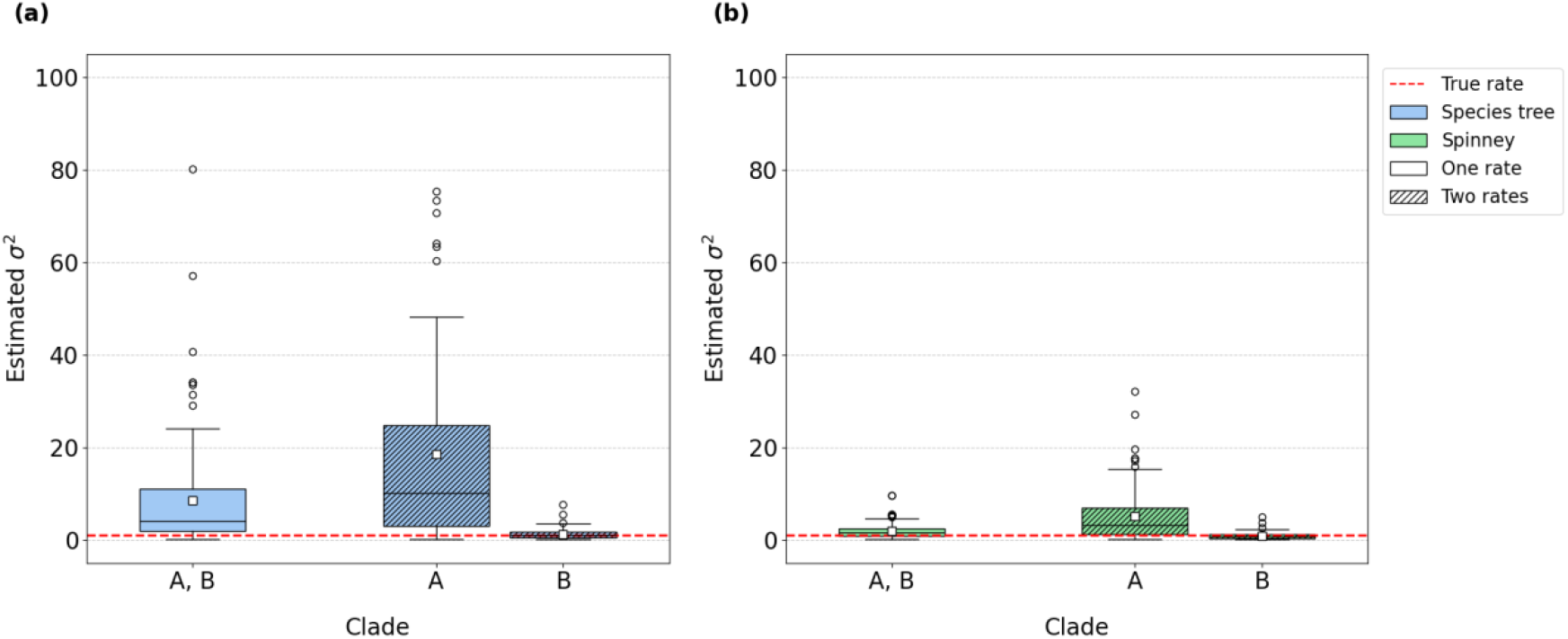
Rate estimates for the six-taxon case with only one rate simulated. Rates are inferred with (a) the species tree or (b) Spinney, with a one-rate model (plain boxes) or two-rate model (shaded boxes). Summary statistics are provided in Supplementary Table 3.

To test whether these overestimates would falsely lead to inferences of rate variation, we recorded the number of cases where the two-rate model was incorrectly favored over the one-rate model via a likelihood ratio test. We then compared these results across different values of *σ*^2^ with 100 replicates per condition (Figure 6). For *σ*^2^ = 0.1, inferences using the species tree incorrectly favored the two-rate model in 31 out of 100 replicates (i.e. a false positive rate of 31%), while Spinney did so in only 14 cases. The pattern was more pronounced at *σ*^2^ = 1, where the species tree inference resulted in 48 incorrect selections (FPR=48%), compared to 15 for Spinney (FPR=15%). At *σ*^2^ = 10, the species tree inference incorrectly favored the two-rate model in 33 cases, whereas Spinney did so in just 12 cases. These results suggest that Spinney effectively reduces the false positive rate by largely accounting for discordant histories.

**Figure 6.**
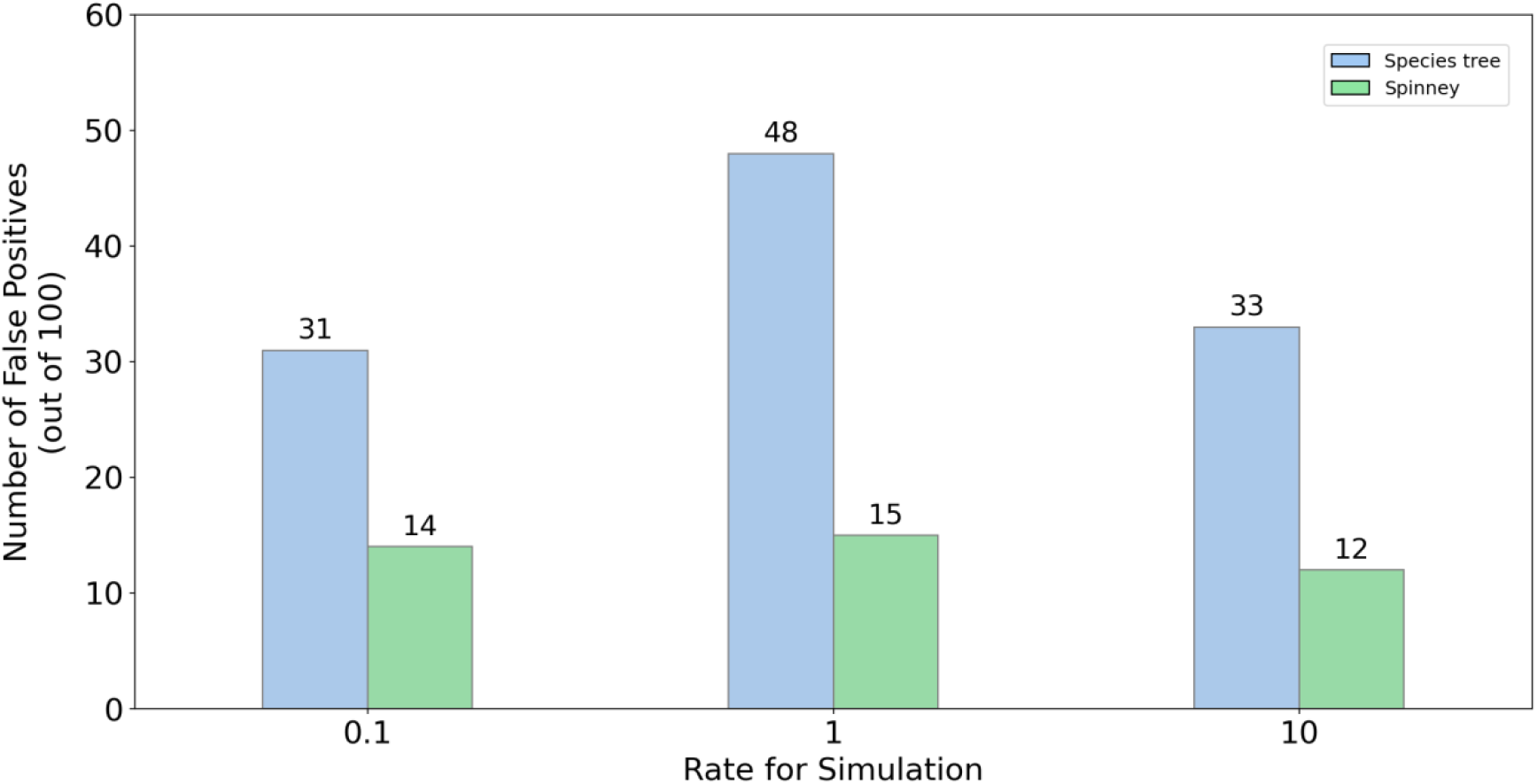
Number of cases in which the two-rate model was incorrectly selected by the LRT when one rate was simulated. The single simulated rate varies along the x-axis, and inference was performed for each condition using both the species tree and Spinney. Spinney reduces the FPR in all conditions, but does not eliminate it entirely.

The results above show that Spinney reduces the false positive rate, but it may do so simply by reducing the overall power of the LRT. To examine the power of both methods when there truly are two rates, we simulated a case in which clade A has a higher rate (i.e. 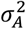 is higher) and also has 25% discordance. Both Spinney and the species tree have similar true positive rates (TPRs) for the two-rate model (Supplementary Table 4). When clade A differed by 100X from clade B, TPRs were more than 90% for both methods. When clade A differed by 10X from clade B, TPRs dropped to 40%. We observed similar patterns when 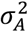 was less than 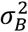 (Supplementary Table 4).

### Ancestral state reconstruction

We evaluated ancestral states across time, focusing on speciation event *T*_2_, which splits taxon *C* from taxa *A* and *B* (Figure 3). Although there is only a single species tree topology, ILS generates four possible genealogical histories at *T*_2_ (corresponding to the expected topologies in Figure 4a).

Although the 95% CIs of ancestral states at time *T*_2_ increased for both methods as ILS increased, inferences using Spinney produced relatively narrower values (Figure 7a). For low discordance, the mean CI widths were 4.11 with a negligible difference from the species tree. For medium discordance, the means were 6.84 and 6.38 for the species tree and Spinney, respectively. For high discordance, the means were 12.82 and 7.40 for species tree and Spinney, respectively (*P*=8.9×10^-18^; Wilcoxon test). The main cause of the larger CIs using the species tree appears to be the higher (incorrect) value of *σ*^2^ used by this method—higher rates mean more spread in the possible ancestral states.

**Figure 7.**
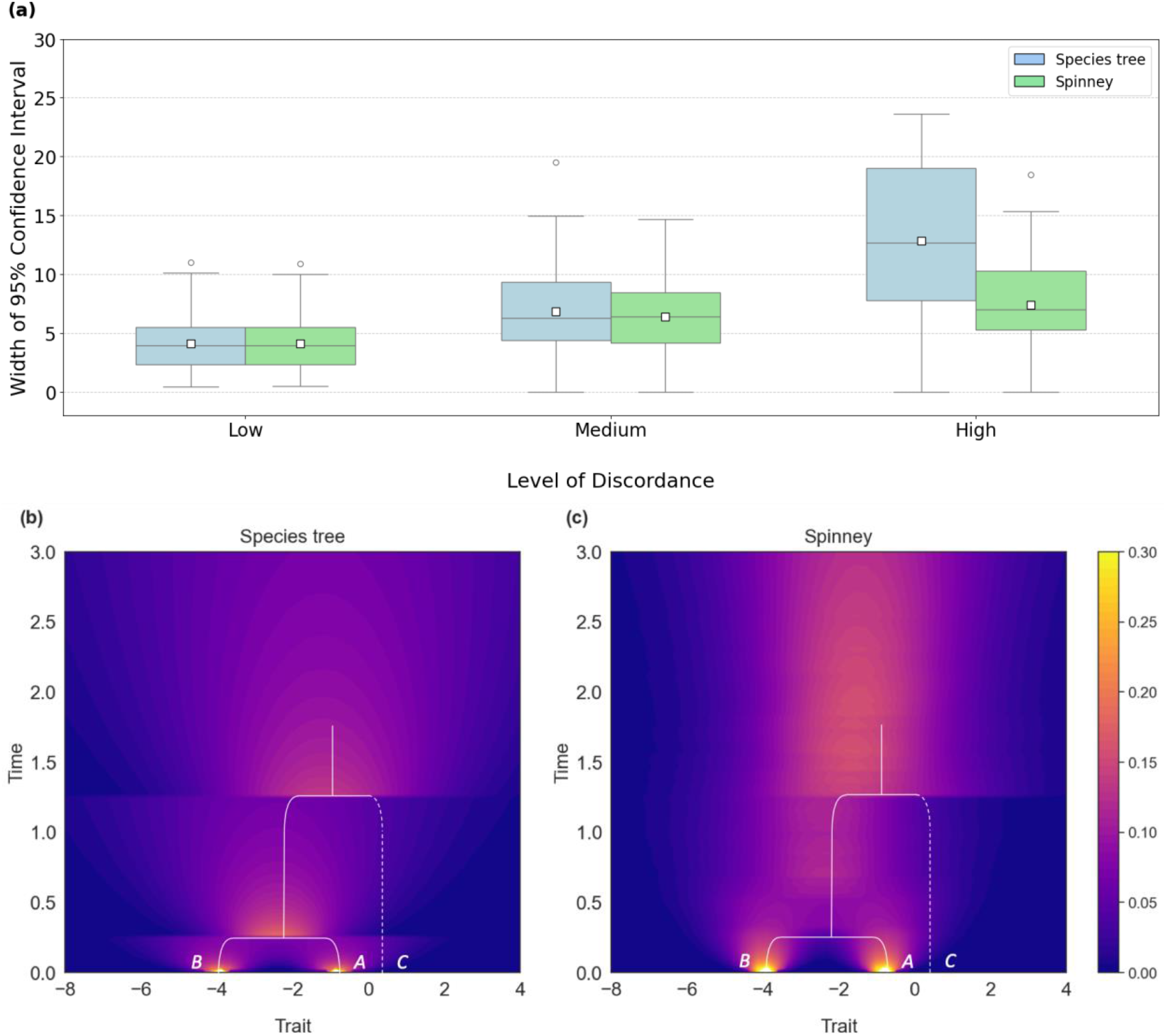
Ancestral states inferred using the species tree and Spinney. (a) The width of the 95% confidence interval at time *T*_2_. (b)-(c) Ancestral state maps of branches extending from taxa *A* and *B* in the species tree. Note that ancestral states for species *C* are not shown, which is why a dashed line is used (species *D* is omitted completely). Ancestral states were inferred with (b) the species tree and (c) Spinney. Colors encode the probability of the ancestral trait value in a specific time bin. The species trees are embedded inside the maps for clarity. Speciation times are *T*_1_=0.25 and *T*_2_=1.25. Species *C* contributes to ancestral states only for *t* >1.25. The trait was simulated with 25% (medium) discordance. Trait values at the tips for taxa *A, B, C*, and *D* were −0.82, −3.95, 0.11, and −3.50, respectively.

We also compared species-tree and Spinney reconstructions of ancestral states across the whole species history, using a trait simulated under medium discordance (Figure 7b, 7c). For 0 ≤ *t* < *T*_1_, both methods behave similarly: the bright regions represent observed trait values that gradually spread under Brownian motion. At *T*_1_, the species tree assumes that species *A* and *B* merge instantly, producing an ancestral state centered between their trait values. In contrast, Spinney allows lineages to coalesce at different times across gene trees, so the spread of ancestral states persists longer into the past. For *T*_1_ < *t* < *T*_2_, the species tree traces only a single, concentrated trajectory, while Spinney aggregates multiple coalescent histories and produces a broader, more uniform distribution of states. At *T*_2_, the species tree again merges species *C* instantaneously, generating a sharp transition with a wider distribution. Beyond *T*_2_, the species tree continues its diffusion process, whereas Spinney maintains a flatter, less concentrated distribution. In general, Spinney accounts for gene tree heterogeneity more correctly and conveys this in inferred ancestral states.

## Discussion

An increasingly common finding in phylogenomics is that a large fraction of gene trees does not match the species tree (e.g. Pollard et al. 2006; White et al. 2009; Jarvis et al. 2014; Copetti et al. 2017; Rouard et al. 2018; Hime et al. 2021; Morales-Briones et al. 2021). Despite the fact that discordant trees are located across the genome and can underlie important traits (e.g. Fontaine et al. 2015; Lamichhaney et al. 2016; Li et al. 2016; Pease et al. 2016; Han et al. 2017; Palesch et al. 2018; Wu et al. 2018; Hibbins et al. 2020; Urban et al. 2021; Feng et al. 2022), PCMs have largely relied on a single, “resolved” species tree to carry out analyses (Hahn and Nakhleh 2016). The use of only a single tree topology when many different topologies exist can cause problems for comparative analyses, including in cases involving continuous traits (Mendes et al. 2018; Bastide et al. 2018; Hibbins and Hahn 2021; Hibbins et al. 2023; Schraiber et al. 2024; Adams et al. 2025).

Here, we have addressed one known issue when using only a single tree in the presence of discordance—that rates of evolution are overestimated overall—and have explored the related, but less-studied problem, of detecting shifts in rates on different parts of a tree. Our approach builds upon the work of Hibbins et al. (2023), who applied the pruning algorithm to calculate trait likelihoods across a set of different tree topologies to account for discordance. While their initial work was limited to a simple three-taxon tree, here we have introduced a program, Spinney, to deal with larger species trees. The results presented above have demonstrated that Spinney reduces, but does not completely remove, the problems of discordance in a principled manner; it should therefore be useful to the research community. Nonetheless, we discuss several caveats, similar approaches, and future directions below.

### The sample of gene trees

For trees with smaller numbers of tips, it is possible to exactly calculate every single relevant gene tree topology (and their weights) to use as input to Spinney. Even when trees have larger numbers of tips, it is really the number of phylogenetic “knots” (cf. Ané et al. 2007) on the tree that determine how many discordant topologies exist. For instance, a species tree with 100 tips might only show 3 topologies if ILS was restricted to a single internal branch. However, with even a bit more discordance, the number of possible gene trees becomes extremely large, requiring that we use only a sample of them as input.

While using more gene trees will always be more precise, this comes at a computational cost because likelihoods must be calculated on each tree (though this can be done in parallel). In our experiments, we used 20 sampled gene trees, which performed well for the species trees we considered, even though the number of possible gene trees was in the millions for some topologies (Figure 4b). Although it may seem surprising in these cases that 20 trees are sufficient, two related aspects of discordance work in our favor. First, many unique tree topologies have very low expected frequencies, and therefore contribute very little to observed trait data. As a result, not using these topologies has little effect on the results. Second, even when there are many phylogenetic knots spread across a species tree, each sampled gene tree provides information about each knot. That is, each gene tree used as input to Spinney can tell us about the dominant concordant and discordant topologies associated with each branch of the species tree, even if each unique tree topology has an individually low probability.

We have suggested two methods that can be used to sample gene trees. The first uses empirical gene trees taken from a dataset. The advantage of such an approach is that one can capture complex causes of discordance in the distribution of trees: for instance, multiple overlapping introgression events. However, input gene trees must be ultrametric and in coalescent units, which is challenging to infer (we have previously found that gene tree inference error has only minor effects; Hibbins et al. 2023). An alternative approach is to infer a species trees whose branch lengths are in coalescent units (e.g. using the approach of Tabatabaee et al. 2023), and then to use this tree to simulate a sample of gene trees. While this will necessarily be a simplified species history, all sampled gene trees can be used directly.

*Testing for multiple rates*. A common use of PCMs is to detect shifts in the rate of evolution along a species tree (O’Meara et al. 2006; Eastman et al. 2011; Rabosky et al. 2014; Revell 2021; Martin et al. 2023). As previous work had already shown that gene tree discordance could drive artifactual patterns of increased evolutionary rates, we reasoned that methods testing for multiple rates might also be misled when levels of discordance differ among clades.

While Hibbins et al. (2023) had to separately analyze parts of a larger tree to make such comparisons, Spinney makes it possible to directly test for multiple rates on a larger tree, while taking into account variable discordance. As shown in our results, Spinney greatly reduced the false positive rate relative to using the species tree alone, but kept the true positive rate on par with the species tree. Nevertheless, Spinney does not completely reduce false positives to their expected level under the null model. We are not sure why this occurs, nor why Spinney overestimates the rate associated with higher discordance, 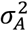, more in this scenario than in the single-rate simulations (see Figure 5b). It may be that something about the shape of the species tree or the number of tips makes it harder to correct for the effect of discordance.

### Similar approaches

There are a few similar approaches to the one used here, both for continuous and discrete traits. For continuous traits, the method of Osmond and Coop (2024) applied to population-level data on ancestral ranges is a clear antecedent. In their original approach, a large number of gene trees from across the genome are used to calculate the dispersal rate and ancestral locations of individuals back in time; this was further extended to the full ancestral recombination graph in Deraje et al. (2025). Osmond and Coop’s approach to calculating dispersal rates is very similar to ours, maximizing the likelihood across trees. Although the resulting estimates are slightly biased, this is likely due to the fact that the underlying model of uniform dispersal does not match actual geographic dispersal routes. Interestingly, for the inference of ancestral locations, Osmond and Coop allow each tree to find its own maximum likelihood value. This highlights a difference in fine-grained population genetic data, where each locus really might have existed in a distinct ancestor living in a distinct geographic location.

A second similar approach, applied to DNA or amino acid sequence data, is the “mixtures across sites and trees” (MAST) model of Wong et al. (2024). In this model, site-likelihoods are calculated under a limited set of alternative tree topologies. The total likelihood of an alignment is maximized over these tree topologies (inferring branch lengths simultaneously) and tree weights. This is very similar to the approach taken by Spinney, with one important distinction: the default setting in MAST also estimates the tree weights from the data. While these weights are not equivalent to gene tree frequencies, they may be highly correlated with these frequencies.

*Future directions*. The approach we take here calculates likelihoods on trees using the pruning algorithm (Felsenstein 1973). Although sometimes overlooked as an approach to PCMs, it has found wide use in a number of programs (e.g. Hahn et al. 2005; FitzJohn 2012; Ho and Ané 2014; Uyeda et al. 2014; Hiscott et al. 2016; Mitov et al. 2020; Bertram et al. 2023; Martin et al. 2023). The pruning algorithm has many advantages over matrix approaches, including the ability to analyze very large trees (e.g. Freckleton 2012), to easily implement a variety of different (non-BM) models of trait evolution (e.g. Boucher and Démery 2016), and to take into account measurement error (e.g. Han et al. 2013). This last feature is extremely important, as measurement error can have drastic consequences for comparative analyses (e.g. Cooper et al. 2016; Beaulieu and O’Meara 2025). In the future, we can imagine combining many of these features with calculations across sets of gene trees—even gene trees possibly inferred in error—to allow for more accurate evolutionary inferences.

## Acknowledgements

We thank Mark Hibbins for very helpful advice early in this project. This research was funded by National Science Foundation grant DBI-2146866.

## Notes

### Competing Interest Statement

The authors have declared no competing interest.

