## Supplementary Figures and Tables for "Estimating the rate of quantitative trait evolution in the presence of gene tree discordance by calculating likelihoods across trees"

### Supplement Information

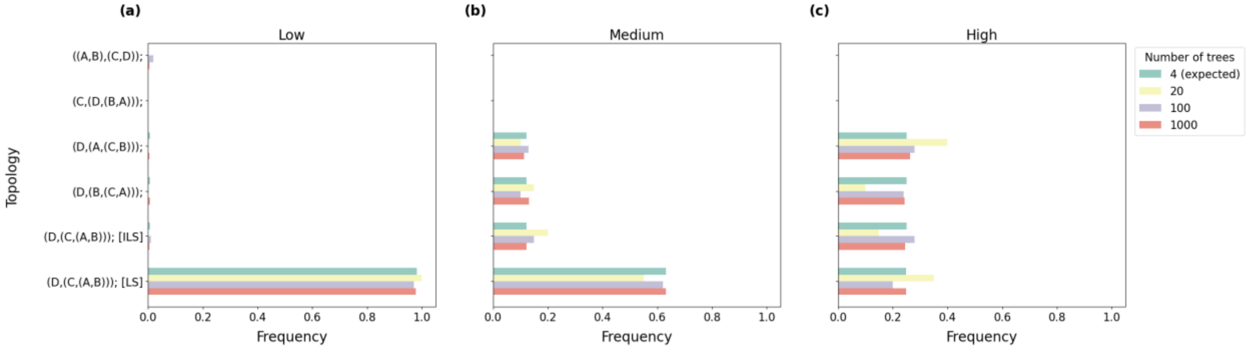


Supplementary Figure 1: Similarity in the frequency distributions of trees for different levels of discordance, with different sample sizes. The species tree used here is (*D*,(*C*,(*A*,*B*))), and simulations with increasing levels of ILS are shown left to right. Green bars are expected gene tree frequencies from multispecies coalescent theory. Yellow bars are gene tree frequencies from the 20 trees used for inference (Figure 1). We also compare to distributions for 100 and 1000 sampled gene trees.

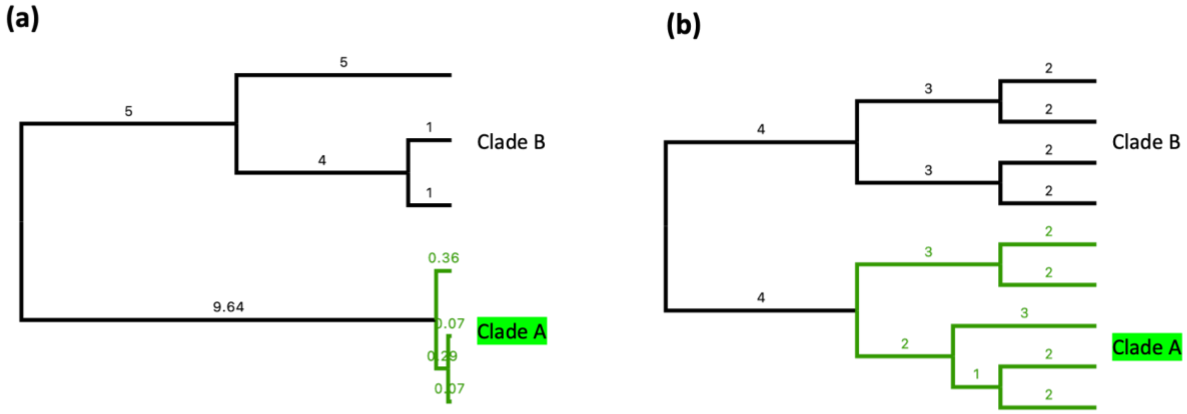


Supplementary Figure 2: Species trees used in two-rate tests, tested for (a) false positive rates and (b) true positive rates. Numbers are branch lengths in coalescent units.

Supplementary Table 1. Estimated $\sigma^2$ for one-rate tests with four-taxon trees (the simulated rate is always $\sigma^2 = 1$)

| Discordance (%) | Species tree | | | Spinney (expected) | | | Spinney (sampled) | | |
|---|---|---|---|---|---|---|---|---|---|
| | Mean | SD | Median | Mean | SD | Median | Mean | SD | Median |
| 1.6 | 1.44 | 1.10 | 1.09 | 1.07 | 0.80 | 0.87 | 1.03 | 0.76 | 0.82 |
| 25 | 3.66 | 3.62 | 2.43 | 1.72 | 1.49 | 1.29 | 1.60 | 1.25 | 1.25 |
| 50 | 12.47 | 15.71 | 7.22 | 2.92 | 2.81 | 1.99 | 3.30 | 2.82 | 2.38 |

Supplementary Table 2. Estimated $\sigma^2$ for one-rate tests with more taxa (the simulated rate is always $\sigma^2 = 1$)

| Number of species | Discordance (%) | Species tree | | | Spinney (sampled) | | |
|---|---|---|---|---|---|---|---|
| | | Mean | SD | Median | Mean | SD | Median |
| 5 | 53 | 1.27 | 0.85 | 1.08 | 1.03 | 0.65 | 0.88 |
| 10 | 68 | 2.00 | 2.30 | 1.57 | 1.11 | 0.57 | 1.00 |
| 20 | 66 | 1.45 | 0.64 | 1.33 | 1.06 | 0.36 | 1.02 |
| 50 | 98 | 1.48 | 0.64 | 1.30 | 1.03 | 0.22 | 1.00 |

Supplementary Table 3. Estimated $\sigma^2$ for the two-clade tree under a one-rate or two-rate model (only one rate with $\sigma^2 = 1$ is simulated)

| Model | Clade | Species tree | | | Spinney (sampled) | | |
|---|---|---|---|---|---|---|---|
| | | Mean | SD | Median | Mean | SD | Median |
| One-rate | A and B | 8.56 | 12.24 | 3.98 | 1.93 | 1.71 | 1.47 |
| Two-rate | A | 18.57 | 24.45 | 10.06 | 5.14 | 5.90 | 3.20 |
| | B | 1.19 | 1.17 | 0.91 | 0.82 | 0.73 | 0.69 |

Supplementary Table 4. True positive rates of the two-clade tree with rate variation.

| $\sigma_A^2$ | $\sigma_B^2$ | Species tree | Spinney |
|---|---|---|---|
| 0.01 | 1 | 97% | 97% |
| 0.1 | 1 | 51% | 54% |
| 1 | 1 | 8% | 7% |
| 10 | 1 | 42% | 42% |
| 100 | 1 | 93% | 91% |
